# Optimization of an improved, time-saving, and scalable, protocol for the extraction of DNA from diverse viromes

**DOI:** 10.1101/2024.06.26.599709

**Authors:** Michael Shamash, Saniya Kapoor, Corinne F. Maurice

## Abstract

**Introduction:** The virome, composed of viruses inhabiting diverse ecosystems, significantly influences microbial community dynamics and host health. The phenol-chloroform DNA extraction protocol for viromes, though effective, is time-intensive and requires the use of multiple toxic chemicals.

**Methods:** This study introduces a streamlined, scalable protocol for DNA extraction using a commercially-available kit as an alternative, assessing its performance against the phenol-chloroform method across human fecal, mouse fecal, and soil samples.

**Results:** No significant differences in virome diversity or community composition were seen between methods. Most viral operational taxonomic units (vOTUs) were common to both methods, with only a small percentage unique to either approach. Alpha- and beta-diversity analyses showed no significant impact of the extraction method on virome composition, confirming the kit’s efficacy and versatility.

**Conclusions:** While the kit approach offers benefits like reduced toxicity and increased throughput, it has limitations such as higher costs and potential issues reliably capturing low-abundance taxa. This protocol provides a viable option for large-scale virome studies, although the phenol-chloroform approach may still be preferable for specific sample types.

## Introduction

The virome, the collection of viruses inhabiting diverse ecosystems, plays important roles in shaping microbial community composition and function^1–5^. In host-associated ecosystems, like the human microbiome, the virome is also a key determinant of host health^6^.

Despite significant advances in the field in recent years, there remain several challenges associated with studying viromes. Many of these challenges are computational, such as identifying novel and highly divergent viruses, or characterizing the host range of uncultured viruses^7^. Others are related to sample processing, especially in low biomass sample types, such as amplifying low amounts of purified virome nucleic acids or dealing with high levels of background contamination (usually of human, mouse, or bacterial origin)^7^.

It is important that virome nucleic acid extraction protocols be efficient and reliable so that we may begin to address some of these analytical challenges. The phenol-chloroform DNA extraction protocol is currently the gold standard for studying viral metagenomes^8–11^. However, this protocol has its limitations, including being time-consuming and requiring the use of multiple toxic chemicals such as phenol and chloroform.

Given the growing interest in studying viromes across diverse environments, including in human^12,13^, animal^14,15^, and soil^16^ communities, there is a need for a faster and scalable extraction protocol which can accommodate increasingly large studies.

In this study, we set out to develop and evaluate a streamlined protocol for virome DNA extractions using a commercially available column-based DNA extraction kit. We compare this kit’s performance to that of the phenol-chloroform protocol to evaluate its efficacy and versatility across a variety of sample types (stool, soil). Overall, we find no significant differences in the resulting sequenced viromes between approaches, with overall sample yield, viral diversity, and community composition being very similar.

## Methods

### Sample collection

Human fecal samples were collected with the approval of protocol A04-M27-15B from the McGill University Institutional Review Board. Participants provided informed written consent for the utilization of their samples and met specific inclusion criteria: they were 18 years of age or older, had no diagnosed gastrointestinal disease, and had not used antibiotics in the 3 months prior to sampling. Fresh fecal samples were collected, aliquoted in an anaerobic chamber, and kept at - 70 °C until processing.

Mouse fecal samples were collected with the approval of animal use protocol MCGL-7999 from McGill University. Fresh fecal samples were collected and kept at -70 °C until processing. All mice had unlimited access to standard chow and water.

Soil samples were collected in sterile 50 mL conical centrifuge tubes from various locations on the downtown campus of McGill University (coordinates: 45.5042 N, 73.5755 W).

### Sample pre-processing

Prior to extracting virome DNA, we first enriched for VLPs in the fecal and soil samples. Samples were resuspended in sterile (0.02µm-filtered) PBS as follows: human fecal samples, ∼200-400mg in 2mL PBS; mouse fecal samples, ∼100mg in 1 mL PBS; soil samples, 10mL soil in 10mL PBS (1:1 volume ratio). Large debris was pelleted by centrifugation at 1,000g for 5 minutes, and the supernatant recovered. Bacterial cells were pelleted by centrifugation at 10,000g for 10 minutes. 500 µL of the VLP-containing supernatant was added to a sterile Ultrafree-MC centrifugal filter unit (MilliporeSigma, Burlington, MA, USA) with 0.22µm pore size and centrifuged at 12,000g for 2 minutes, and 400 µL of purified VLPs was recovered. The VLP suspension was cleaned by addition of 100 µL chloroform (final chloroform concentration: 20% v/v), followed by thorough vortexing and centrifugation at 21,000g for 5 minutes. The VLP-containing supernatant (upper layer) was carefully recovered and transferred to a new tube containing 5 µL (10 U) TURBO DNase (ThermoFisher Scientific, Waltham, MA, USA), 50 µL 10X TURBO DNase buffer, and 1 µL (approx. 125 U) Benzonase DNase (MilliporeSigma, Burlington, MA, USA). The sample was incubated at 37°C with mild shaking for 90 minutes. To stop the DNase digestion reaction, 18 µL of a 500 mM EDTA (pH 8) solution was added, followed by heat inactivation of the enzymes at 75°C for 30 minutes. Sterile PBS was added to bring the samples up to 800 µL in volume: 400 µL for phenol/chloroform extraction, and 400 µL for kit extraction.

### DNA extraction – phenol/chloroform approach

This protocol is adapted from Thurber et al^8^, where 40 uL of 200X TE buffer (2M tris, 200 mM EDTA, pH 8.5), 440 uL formamide, and 10 uL UltraPure glycogen (20mg/mL stock; ThermoFisher Scientific, Waltham, MA, USA) were added to 400 µL purified VLPs and incubated at room temperature for 30 minutes. Two volume equivalents (approx. 1,780µL) of room temperature 100% ethanol were added to the sample, and DNA was pelleted by centrifugation at 10,000g for 20 minutes at 4°C. The supernatant was carefully removed and discarded, and the DNA pellet was washed twice with 1 mL of ice-cold 70% ethanol, centrifuging as above between washes. The final pellet was dried for 5 minutes at room temperature and resuspended in 567 µL of 1X TE buffer (10 mM tris, 1mM EDTA, pH 8). We then added 30 µL of 10% (w/v) SDS and 3 µL of Proteinase K (20 mg/mL stock; ThermoFisher Scientific, Waltham, MA, USA) to the sample and briefly vortexed before incubation at 45°C for 1 hour with gentle shaking. After incubation, 100 µL of 5M NaCl and 80 µL of CTAB buffer (1.1M NaCl, 450 mM CTAB/cetyltrimethylammonium bromide) were added, the sample vortexed and incubated at 65°C for 10 minutes with gentle shaking. One volume equivalent (approx. 780 µL) of chloroform:isoamyl alcohol 24:1 (Sigma-Aldrich) was added and mixed well, transferred to a light phase lock gel tube (PLG tube; Quantabio, Beverly, MA, USA), and centrifuged at 12,000g for 5 minutes. The aqueous (upper) phase was transferred to a new PLG tube to which another 1 volume equivalent of phenol:chloroform:isoamyl alcohol 25:24:1 (pH 8, Invitrogen) was added, mixed well, and centrifuged as above. After transferring the aqueous (upper) phase to a new PLG tube, this step was repeated with 1 volume equivalent of chloroform:isoamyl alcohol 24:1. The aqueous phase was then recovered, added to a tube containing 550 µL (∼0.7 volume equivalents) ice-cold 100% isopropanol, and stored overnight at -20°C for DNA precipitation. The next day, DNA was pelleted by centrifugation at 13,000g for 15 minutes at 4°C. The pellet was washed once with 500 µL ice-cold 70% ethanol and pelleted again as above. The final pellet was air-dried at room temperature, resuspended in 50 µL tris buffer (10 mM Tris-Cl, pH 8), and stored in a DNA LoBind tube (Eppendorf, Hamburg, Germany) at -20°C until library preparation.

### DNA extraction – kit approach

DNA was extracted using the QIAGEN MinElute Virus Spin Kit (QIAGEN, Hilden, Germany) according to the manufacturer’s instructions, with the following modifications to incorporate a larger sample input volume. Fifty µL QIAGEN Protease and 400 µL Buffer AL were added to 400 µL purified VLPs, vortexed, and incubated at 56°C for 15 minutes. We then added 500 µL 100% ethanol to the sample, vortexed, and half of the sample (∼675 µL) was added onto a QIAamp MinElute column, and centrifuged at 6,000g for 1 minute. The filtrate was discarded and the remaining sample was added to the same QIAamp MinElute column, centrifuged as above. The remainder of the protocol remained unchanged from the manufacturer’s instructions. DNA was eluted from the column using 50 uL Buffer EB (tris buffer; 10 mM Tris-Cl, pH 8; 5-minute incubation before elution), and stored in a DNA LoBind tube (Eppendorf, Hamburg, Germany) at -20°C until library preparation. The included Carrier RNA was not used for any of the DNA extractions.

### Library preparation and sequencing

Extracted vDNA from both procedures was quantified using the Qubit 1X dsDNA High Sensitivity kit (Invitrogen). Sequencing libraries were prepared using the Illumina DNA Prep kit (Illumina, San Diego, CA, USA) according to the manufacturer’s instructions, maximizing sample input (30 µL) and number of PCR cycles (12 cycles). The final libraries were quantified using the Qubit 1X dsDNA High Sensitivity kit and the Bioanalyzer High Sensitivity DNA kit (Agilent Technologies, Santa Clara, CA, USA). An equimolar pool of libraries was created and sequenced on an Illumina MiSeq instrument with 150bp paired-end reads (SeqCenter, Pittsburgh, PA, USA).

### Bioinformatic analysis of sequence data

Raw reads were trimmed and filtered with fastp (v0.20.1)^17^ using the following criteria: -- detect_adapter_for_pe -q 15 --cut_right --cut_window_size 4 --cut_mean_quality 20 -- length_required 100. Trimmed reads were decontaminated for human and mouse genomic DNA with bowtie2 (v2.4.2)^18^ using the *Homo sapiens* GRCh38 and *Mus musculus* GRCm39 references, respectively. metaSPAdes (v3.15.4)^19^ was used to conduct *de novo* assembly of each sample individually using default settings. VIBRANT (v1.2.1)^20^ was used on these filtered contigs to identify viral sequences with the following settings: 1kb minimum length, virome mode. Viral contigs were dereplicated into viral operational taxonomic units (vOTUs) using BLASTN (v2.14.0)^21^ followed by the anicalc.py and aniclust.py scripts from CheckV^22^ with the following parameters: 95% average nucleotide identity (ANI) over 85% of the contig’s length. Bowtie2 (v2.4.2)^18^ was used to map decontaminated trimmed/filtered reads to the resulting dereplicated set of vOTUs, and a coverage summary report was generated with Samtools (v1.13)^23^ using the ‘samtools coverage’ command. R (v4.2.2) was used for all diversity analyses with the following packages: phyloseq (v1.42.0)^24^ and vegan (v2.6.4)^25^. Flextable (v0.8.5)^26^ was used to generate the table in Figure 2C.

## Code and data availability

Code used for data analysis is available at: https://github.com/mshamash/vdna_protocol_manuscript. Virome sequencing reads (human and mouse sequences removed) are available on the NCBI SRA using accession number PRJNA1125394.

## Results

To compare our new viral DNA protocol (KIT) with the previous standard, phenol-chloroform extractions (PC), we collected samples from a variety of environments which were each processed with both extraction methods: 2 soil samples, 7 human fecal samples, and 4 mouse fecal samples (**Figure 1**). DNA yields were not significantly different across protocols for soil (42 ± 2 pg/µL KIT, 47 ± 32 pg/µL PC), human fecal (249 ± 141 pg/µL KIT, 536 ± 314 pg/µL PC) and mouse fecal (45 ± 14 pg/µL KIT, 126 ± 42 pg/µL PC) samples (p > 0.05, Wilcoxon signed-rank test).

**Figure 1.**
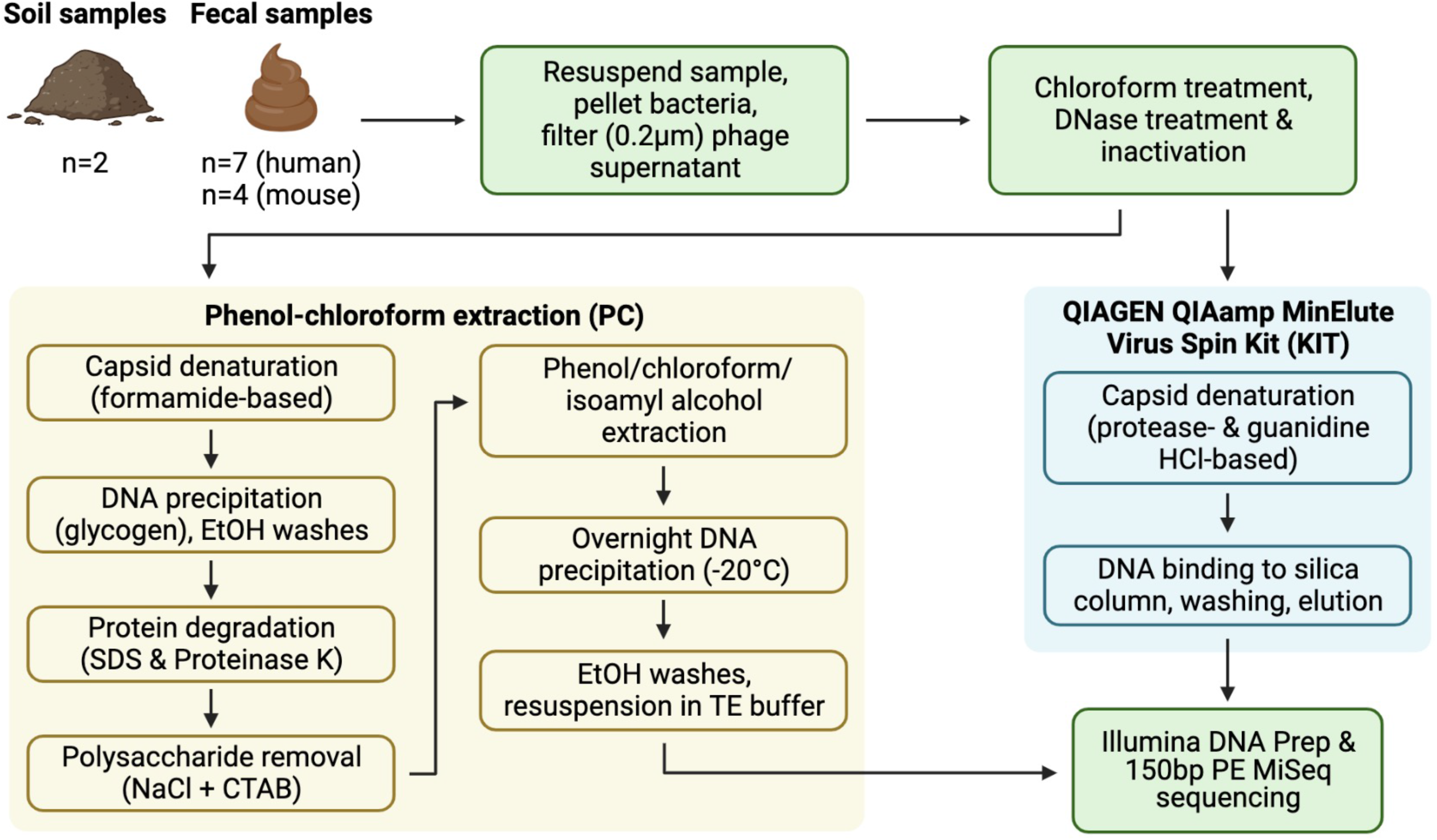
Overview of sample processing pipeline. All samples underwent resuspension in sterile PBS, filtration to remove debris and bacterial or eukaryotic cells, chloroform treatment, and DNase treatment (with subsequent DNase inactivation). Samples were then split to be processed with both the phenol-chloroform extraction (PC) and QIAGEN QIAamp MinElute Virus Spin Kit (KIT) DNA extraction protocols. DNA libraries were prepared with the Illumina DNA Prep kit and sequenced on an Illumina MiSeq instrument with 150bp paired-end reads.

**Figure 2.**
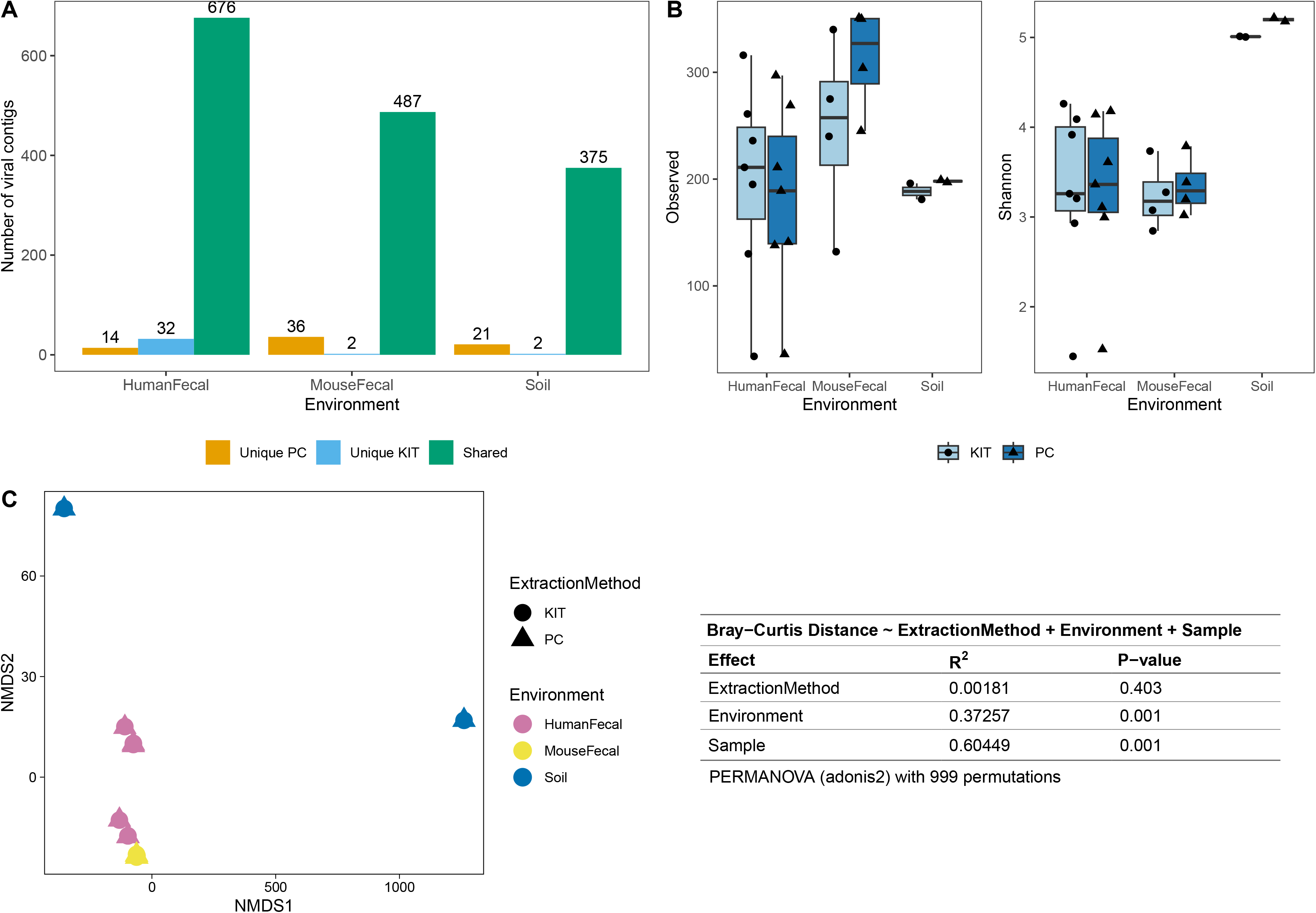
PC and KIT extraction methods result in equivalent virome communities. **(A)** The number of vOTUs detected only within the phenol-chloroform (PC) or kit (KIT) extraction methods, or shared between both methods. **(B)** Observed richness (left) and Shannon diversity (right) of viromes according to environment and extraction method. No statistical differences were observed between extraction methods within any environment (Wilcoxon signed-rank test, *p* > 0.05). **(C)** Non-metric multidimensional scaling (NMDS) of Bray-Curtis distances of vOTUs between samples, colors representing sample environment and shapes representing extraction method (NMDS stress = 9.5 x 10^−5^). Tests for significant differences in Bray-Curtis dissimilarity were conducted using PERMANOVA (adonis2) with 999 permutations, and summary statistics are reported in the table.

After sequencing, assembly, and viral detection, a total of 722, 525, and 398 vOTUs were detected in human fecal, mouse fecal, and soil samples, respectively. Of these, most vOTUs were common to both PC and KIT viromes: 676 (94%) of all vOTUs in human fecal viromes, 487 (93%) of all vOTUs in mouse fecal viromes, and 375 (94%) of all vOTUs in soil viromes (**Figure 2A**). Few contigs (6.4% of human fecal, 7.2% of mouse fecal, and 5.8% of soil vOTUs) were detected exclusively in either PC or KIT viromes (**Figure 2A**).

We next characterized the alpha- and beta-diversity of our samples, to evaluate compositional differences in viromes which may be due to extraction method. While alpha-diversity, at the observed richness and Shannon diversity levels, was numerically higher with the PC extraction method (mean 206, 229, 418 observed vOTUs, and 2.99, 2.28, 5.78 Shannon index for human fecal, mouse fecal, and soil viromes, respectively, with KIT; and mean 189, 278, 554 observed vOTUs, and 2.95, 2.39, 6.18 Shannon index for human fecal, mouse fecal, and soil viromes, respectively, with PC), these differences were not significant (**Figure 2B**). Pairwise Bray-Curtis distances were calculated between all samples and plotted on an NMDS plot (**Figure 2C**). For each sample, PC and KIT viromes clustered closely together regardless of the environment. A PERMANOVA analysis confirmed that the distances between samples was explained primarily by environment (R2 = 0.373, *p* = 0.001) and the sample itself (R2 = 0.605, *p* = 0.001), rather than the extraction method (R2 = 0.002, *p* = 0.403; **Figure 2C**).

## Discussion

In this study, we describe a streamlined protocol for virome DNA extractions using a commercially available column-based DNA extraction kit, with some upstream steps. This new approach has several advantages over the gold standard phenol-chloroform protocol, including reduced dependency on toxic chemicals, increased throughput, and improved ease-of-use.

We compared this kit’s performance to that of the phenol-chloroform protocol to evaluate its relative performance across a variety of sample types, including human fecal, mouse fecal, and soil samples (**Figure 1**). The effect of the kit used did not significantly affect virome diversity or community composition (**Figure 2**).

Most of the assembled vOTUs were detected in both KIT and PC viromes, with few vOTUs unique to either approach (**Figure 2A**). The presence of unique vOTUs to either KIT or PC viromes did not have a significant effect on community alpha- or beta-diversity (**Figures 2B, 2C**). The unique vOTUs were low in abundance (mean unique vOTU abundance of 0.117% ± 0.065%), indicating that both extraction methods were equally good at capturing high-abundance vOTUs.

Despite its advantages, the kit approach has a few limitations. This kit still includes a chloroform step upstream of extraction which destroys the membrane of enveloped viruses^27^. Cost may be another factor, as kits are generally more expensive than the different reagents used in the phenol-chloroform method. Furthermore, reliance on proprietary reagents may be an issue if the manufacturer changes their formulation. Finally, while kits offer high consistency, they may not always result in the highest yield for all sample types. Researchers specifically targeting low-abundance virome members may still wish to use the phenol-chloroform method. For researchers working with low biomass samples, it is important to test and validate the kit’s performance with this sample type.

## Funding information

This work was supported by a Canadian Institutes of Health Research Canada Project grant (PJT-175065) and an NSERC Discovery grant (RGPIN 2023-04216) to CFM. CFM is a Tier 2 Canada Research Chair in gut microbial interactions. MS is supported by the Canadian Institutes of Health Research Canada Graduate Scholarship to Honor Nelson Mandela (CIHR CGS-D; #DF2-187718), and the Fonds de recherche du Québec-Santé: Bourse de formation au doctorat (FRQS; #311071).

## Notes

**Competing interests statement** The authors have no conflicts of interest to disclose.

https://github.com/mshamash/vdna_protocol_manuscript

### Competing Interest Statement

The authors have declared no competing interest.

